# Evidence for close molecular proximity between reverting and undifferentiated cells

**DOI:** 10.1101/2022.02.01.478637

**Authors:** Souad Zreika, Camille Fourneaux, Elodie Vallin, Laurent Modolo, Rémi Seraphin, Alice Moussy, Elias Ventre, Matteo Bouvier, Anthony Ozier-Lafontaine, Arnaud Bonnaffoux, Franck Picard, Olivier Gandrillon, Sandrine Giraud

**Affiliations:** Laboratory of Biology and Modelling of the Cell, Université de Lyon, Ecole Normale Supérieure de Lyon, CNRS, UMR5239, Université Claude Bernard Lyon 1, Lyon, France; Azm Center for Research in Biotechnology and its Applications, LBA3B, EDST, Lebanese University, Tripoli 1300, Lebanon; Ecole Pratique des Hautes Etudes, PSL Research University, UMRS951, INSERM, Univ-Evry, Paris, France; Inria Team Dracula, Inria Center Grenoble Rhone-Alpes, Grenoble, France; Institut Camille Jordan, CNRS UMR 5208, Université Claude Bernard Lyon 1, Villeurbanne, France; Vidium solutions, Lyon, France; Nantes Université, Centrale Nantes, Laboratoire de mathématiques Jean Leray, LMJL, F-44000 Nantes, France

## Abstract

According to Waddington’s epigenetic landscape concept, the differentiation process can be illustrated by a cell akin to a ball rolling down from the top of a hill (proliferation state) and crossing furrows before stopping in basins or “attractor states” to reach its stable differentiated state. However, it is now clear that some committed cells can retain a certain degree of plasticity and reacquire phenotypical characteristics of a more pluripotent cell state. In line with this dynamic model, we have previously shown that differentiating cells (chicken erythrocytic progenitors (T2EC)) retain for 24 hours the ability to self-renew when transferred back in self-renewal conditions. Despite those intriguing and promising results, the underlying molecular state of those “reverting” cells remains unexplored. The aim of the present study was therefore to molecularly characterize the T2EC reversion process by combining advanced statistical tools to make the most of single cell transcriptomic data. For this purpose, T2EC, initially maintained in a self-renewal medium (0H), were induced to differentiate for 24h (24H differentiating cells); then a part of these cells was transferred back to the self-renewal medium (48H reverting cells) and the other part was maintained in the differentiation medium for another 24h (48H differentiating cells). For each time point, cell transcriptomes were generated using scRT-qPCR and scRNAseq. Our results showed a strong overlap between 0H and 48H reverting cells when applying dimensional reduction. Moreover, the statistical comparison of cell distributions and differential expression analysis indicated no significant differences between these two cell groups. Interestingly, gene pattern distributions highlighted that, while 48H reverting cells have gene expression pattern more similar to 0H cells, they retained traces of their engagement in the differentiation process. Finally, Sparse PLS analysis showed that only the expression of 3 genes discriminates 48H reverting and 0H cells. Altogether, we show that reverting cells return to an earlier molecular state almost identical to undifferentiated cells and demonstrate a previously undocumented physiological and molecular plasticity during the differentiation process, which most likely results from the dynamic behavior of the underlying molecular network.

## Introduction

The integration and processing of endogenous and exogenous information constitute a fundamental requirement for cells to ensure functions and survival of unicellular or multicellular organisms. Cellular decision-making is then at the core of the physiological or pathological functioning of living organisms. Early views of the mechanisms governing cell-fate decision-making, and in particular cell differentiation, were based on bulk population data, leading to an over-simplifying deterministic framework. In these first views, cell commitment to a predefined cell-type was thought to be triggered through a stereotyped sequence of intermediate states under the influence of specific signals (1).

Single-cell approaches have allowed to change the scale of observation of many molecular processes and revealed that an important heterogeneity in gene expression lies at the heart of isogenic cell populations (2,3). Stochasticity in gene expression arises from different causes, such as the probabilistic nature of molecular interactions or transcriptional bursts (4). Cell-to-cell variability is visible at all omics levels of gene expression, but is being widely studied at the transcriptomic level since various molecular biology tools are available for this scale of investigation (5). Overall, this heterogeneity in gene expression has been shown to be critical for the process of differentiation, as it provides diversity without the cost of hardwire-encoded fate programs (6,7).

Furthermore, single-cell studies have also enabled the development of stochastic models to describe differentiation from single-cell transcriptomics data. One of the best-known stochastic model is Conrad Waddington’s landscape, that also includes the non-genetic part of cell-to-cell heterogeneities (8). According to Waddington’s model, the shape of the landscape is determined by Gene Regulatory Networks (GRN) and state transitions are modelled as channelling events: a cell, presented as a ball, starts from a mountain top and crosses valleys before reaching stable state by occupying basins or attractor states, shaped by an underlining GRN (9). Once this stable state is reached, the state potential decreases and the associated cell-fate is restricted or even irreversible (10). However, it is now clearly accepted that some cells retain fate plasticity (11,12). Under the forced modification of transcription factors stoichiometry, a cell that have reached a differentiated state can return to a more pluripotent stage challenging the classical hierarchical view of differentiation (13,14). Quite interestingly, spontaneous fate reversion can be observed under physiological or damaging condition where progenitors or even more committed cells return to an earlier stage, potentially more pluripotent and re-acquire progenitor or stem-cell-like phenotypes and characteristics (15–18). In this view, our recent study has shown that chicken primary erythroid progenitor cells (T2EC) have retained the capacity to go back to self-renewal state for up to 24H after the induction of differentiation before they irreversibly engaged in the differentiation process (19). Despite intriguing and promising results, the molecular determinants of this so-called fate reversion and the molecular characterization of the reverting cells remain unexplored.

In this work, we go beyond the cellular and phenotypic characterization of the cell reversion process. We characterize the gene expression of primary erythroid progenitors and question if reverting cells undergo an actual fate reversion i.e. in addition to regaining a comparable cellular state, reacquire a molecular state similar to undifferentiated cells.

For this, differentiation of self-renewing cells was induced by medium change during 24H. Then we splitted the differentiating population so that half could pursue differentiation, and the second half was shifted back in self-renewal medium (FIGURE 1). To provide robust quantitative measurements of gene expression variability, we combined a highly sensitive targeted quantification method (scRT-qPCR) with genomewide scRNASeq data to characterize the transcriptome of each population at single cell level: undifferentiated (0H), differentiating (24H and 48H) and reverting (48H reverting) cells. Our statistical analyses show that 48H reverting cells and undifferentiated cells were much more similar, whereas a separation was clearly visible between cells maintained in differentiation (48H differentiating cells) and cells in reversion (48H reverting cells). Furthermore, statistical comparison of cell distributions indicated no significant differences between 0H cells and 48H reverting cells. Moreover, gene expression pattern distribution of 48H reverting cells showed a shift towards expression pattern distribution of 0H cells. Finally, we identified genes that discriminate 48H reverting cells and 0H cells. Using sparse Partial Least Square (20), we were able to show that the expression of 3 genes, *HBBA, TBC1D7* and *HSP90AA1*, was discriminant between 48H reverting cells and 0H cells showing that reverting cells kept transcriptional traces of their induction to differentiation. In conclusion, our results show that reverting cells display gene expression patterns that are very similar to undifferentiated cells while retaining traces of their response to differentiation induction, which suggests an almost complete molecular reversion after 24H of differentiation induction.

**Figure 1:**
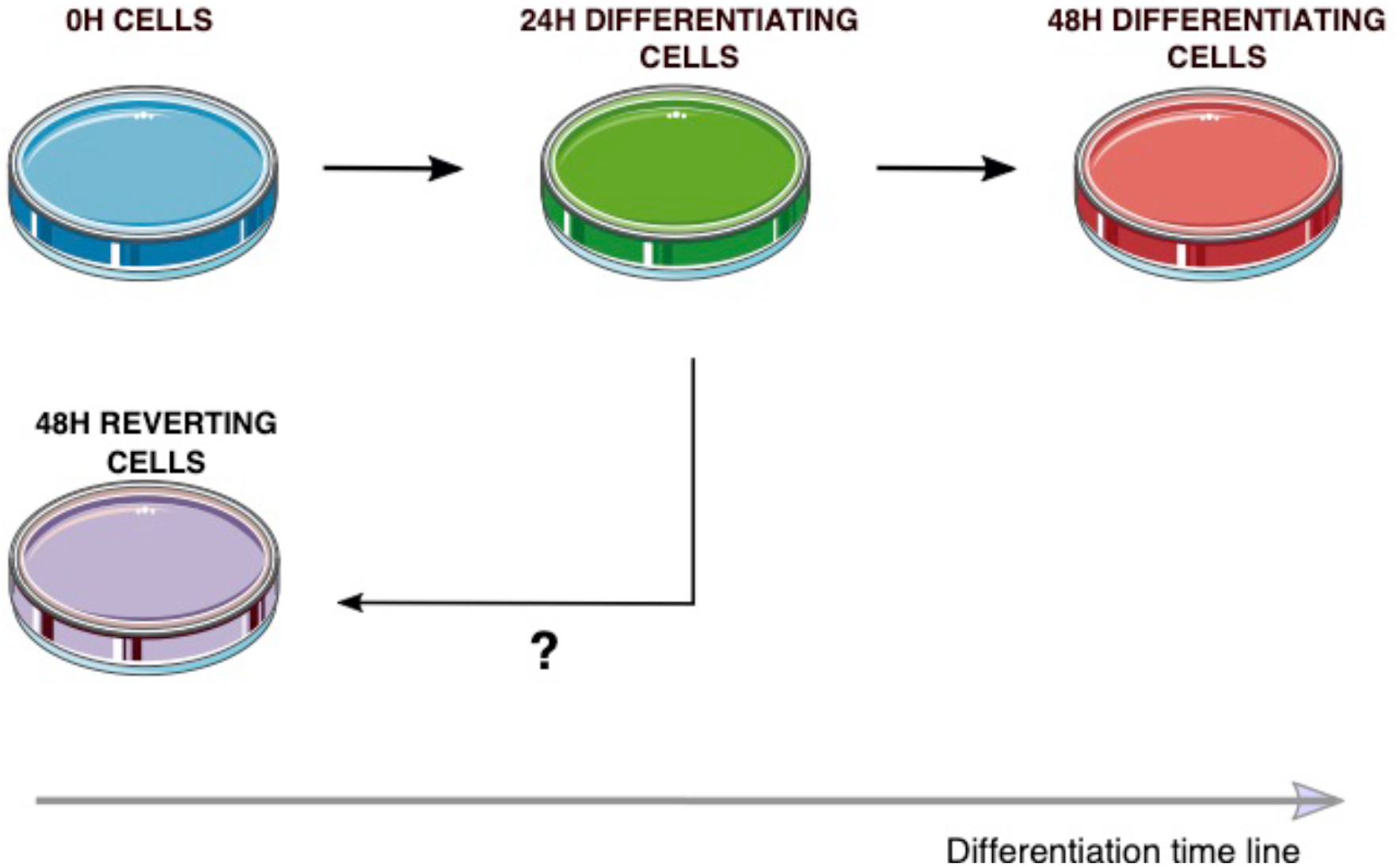
Experimental design. At 0H, cells grown in self-renewal medium are shifted in differentiation medium. 24H later, the cell population is divided in two, half being kept in differentiation medium and half being grown back into self-renewal medium. At each time point, 192 cells are collected for each subsequent experiment: scRT-qPCR and scRNAseq.

## Results

### Robustness of single-cell transcriptomics analysis

We sought to characterize at the molecular level the cells that were induced to differentiate for 24 hours and that retained the ability to proliferate when placed back into self-renewal medium. We used two different complementary single-cell transcriptomic technologies, scRT-qPCR and scRNAseq. scRT-qPCR allows for highly sensitive quantification but is knowledge-driven and offers information of a limited number a genes while scRNAseq, although less precise for low expression level (21), enables genome-wide quantification without prior knowledge. Furthermore, using two different single-cell technologies allowed us to cross validate our observations and point toward robust conclusions.

We first obtained by scRT-qPCR the expression level of 83 genes involved in T2EC differentiation, in 173, 173, 168, 171 cells for 0H, 24H, 48H of differentiation and 48H reverting cells, respectively. Those genes are known to distinguish cells along the differentiation process and include sterol biosynthesis, metabolism, globin subunits, and transcription factors expressed by erythroid progenitors as published in (19). The robustness of our measurements was confirmed by a Pearson’s correlation of 0,85 (p-value = 2.2e-16) between our experiments and the published data (19). To investigate fate-reversion genome-wide by scRNASeq, we adapted the MARSseq protocol ((22) - see material and methods). Then we obtained gene expression levels in 174, 181, 169, 186 single-cells for 0H, 24H, 48H of differentiation and 48H reverting cells, respectively. The concordance between scRT-qPCR and scRNAseq data was confirmed by a Pearson’s correlation of 0,73 (p-value = 1,34e-13) between the 74 genes common to both datasets.

### Similarity between reverting and undifferentiated cells revealed by dimension reduction

We used UMAP to uncover potential similarities between 48H reverting cells and subgroups of differentiating cells by projecting the 4 conditions (FIGURE 2: Panel A, scRT-qPCR data and Panel B, scRNAseq data). Then we focused on the normal differentiation process using the 3 times points of differentiation (0H, 24H and 48H differentiating cells) (FIGURE 2: Panel C - H). For both experiments, pairwise representations show that 24H differentiating cells tend to overlap with both 0H cells (Figure 2: Panel C and D) and 48H differentiating cells (Figure 2: Panel E and F). On the contrary, the undifferentiated cells and 48h differentiating cells clearly differ. Interestingly, pairwise representations also reveal that 48H reverting cells separate well from the 48H differentiating cells (FIGURE 2: I and J) and from 24H cells (FIGURE 2: K and L), but are visually not distinguishable from the 0H cells (FIGURE 2: M AND N). Almost identical results were observed when, instead of plotting cells on the UMAPs calculated from the mix of the 4 conditions, we recalculated the UMAPs for each pair of conditions (Supplementary 1). Those analyses suggest that the transcriptomes of 48H reverting cells are more similar to the undifferentiated cells than to any other condition at both scales of observation. This was further confirmed by the pairwise statistical comparison of average scRNAseq distributions ((23) - see material and methods). As shown in Table 1, the average transcriptomes of 48H reverting and 48H differentiating cells are significantly different, as well as of undifferentiated and 48H differentiating cells. In contrast, no significant difference in average transcriptomes was detected between 0H and 48H reverting conditions (p-value >> 0.05), indicating a very close proximity of 48H reverting cells to undifferentiated cells.

**Figure 2:**
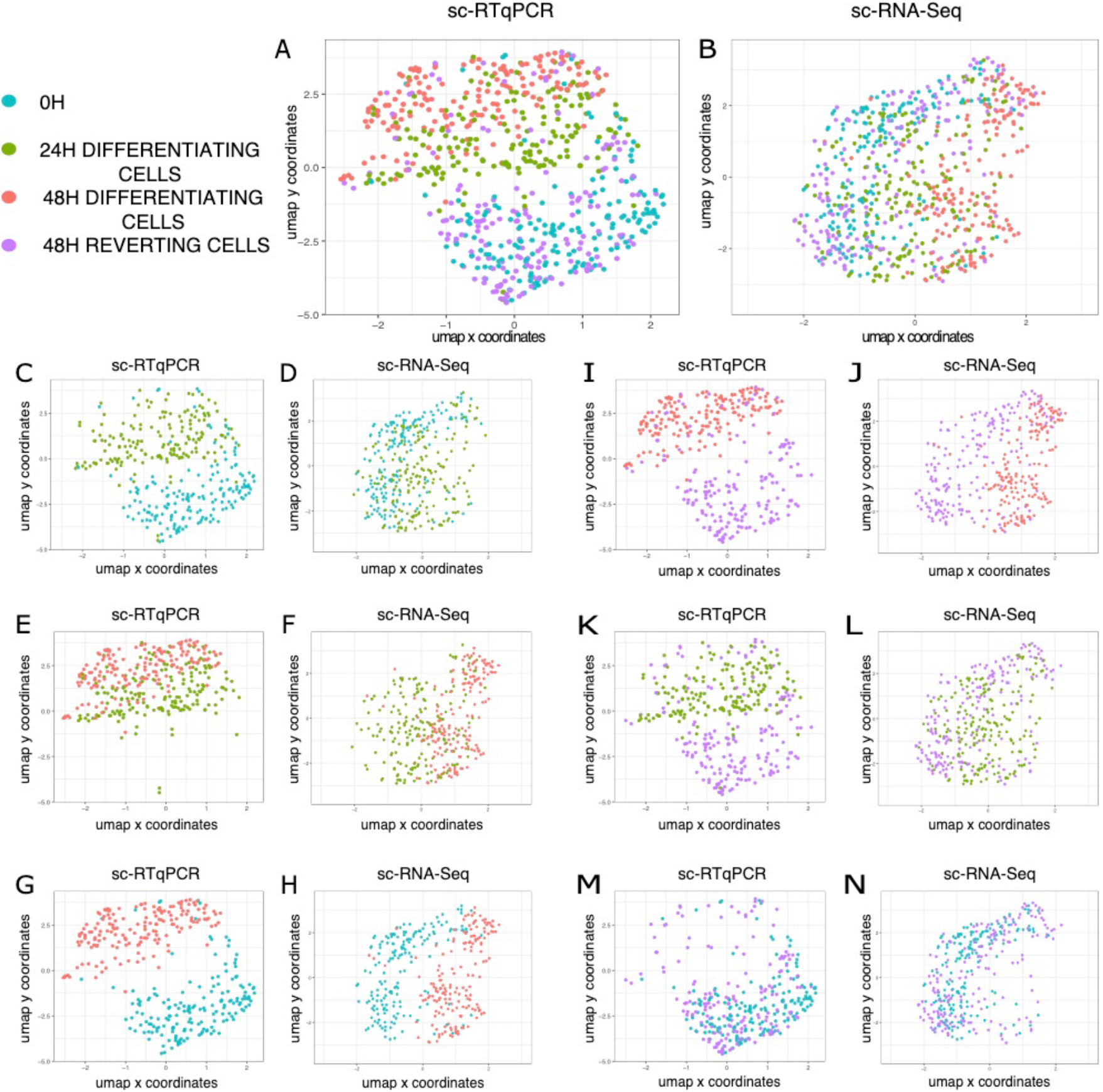
UMAP visualization of scRT-qPCR and scRNAseq data. All UMAPs are calculated using the 4 biological conditions. 0H cells are displayed in blue, 24H cells in green, 48H differentiating cells in red and 48H reverting cells in purple. Panels A, C, E, G, I, K and M: scRT-qPCR data Panels B, D, F, H, J, L and N: scRNAseq data Panels A and B: all 4 conditions Panels C and D: 0H and 24H differentiating cells Panels E and F: 24H and 48H differentiating cells Panels G and H: 0H and 48H differentiating cells Panels I and J: 48H differentiating and 48H reverting cells Panels K and L: 24H and 48H reverting cells Panels M and N: 0H and 48H reverting cells

### 48H reverting cells and undifferentiated cells have similar gene expression patterns

We then questioned if 48H reverting cells had gene expression patterns identical to 0H cells or retained, for some genes, an expression pattern more similar to 24H or 48H differentiating cells.

Pairwise scRNAseq DE analysis revealed that the “normal” erythrocyte differentiation process showed an increase in the expression of hemoglobin related genes during the kinetics (Hemoglobin subunit epsilon 1 (*HBBA*), Hemoglobin Alpha-Locus 1 (*HBA1*), and Hemoglobin Alpha, subunit D (*HBAD*)) (FIGURE 3: panel A, B and C). On the other hand, 0H cells expressed high level of *LDHA* (Lactate Dehydrogenase A), marker for glycolysis metabolism used by self-renewing cells (24) and *ID2* (Inhibitor Of DNA Binding 2) coding for a transcription factor involved in differentiation inhibition (25).

**Figure 3:**
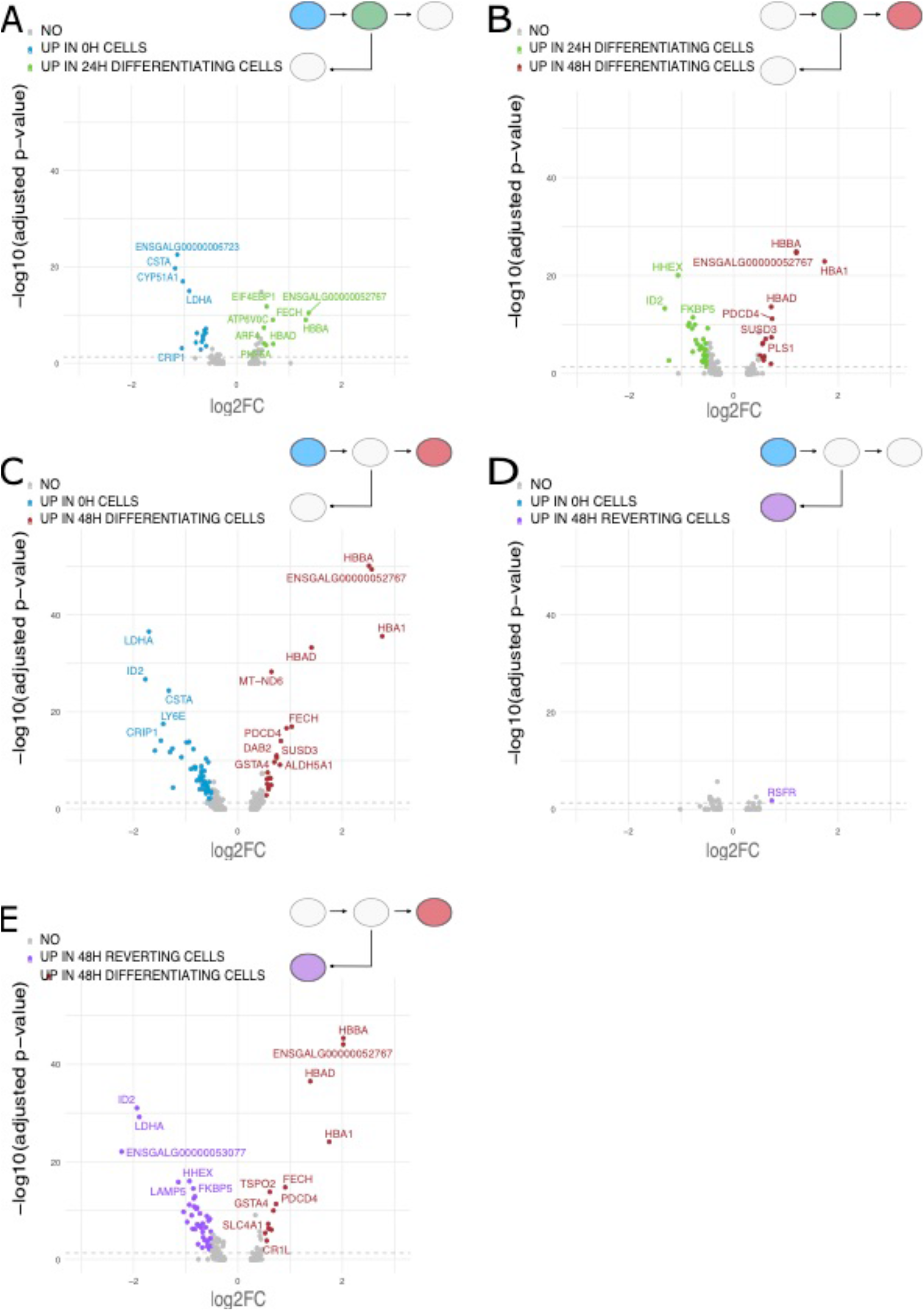
Volcano plot of genes from scRNAseq data differentially expressed between conditions analyzed two by two. Genes are considered significantly differentially expressed when the fold change is equal or above 0,5 and adjusted p-value is below 0,05 (grey dotted line). Panel A: 0H and 24H differentiating cells. Blue dots represent significantly up-regulated genes in 0H condition and green dots represent significantly up-regulated genes in 24H condition. Panel B: 24H differentiating and 48H differentiating cells. Green dots represent significantly up-regulated genes in 24H differentiating cells and red dots represent significantly up-regulated genes in 48H differentiating cells. Panel C: 0H and 48H differentiating cells. Blue dots represent significantly up-regulated genes in 0H cells and red dots represent significantly up-regulated genes in 48H differentiating cells. Panel D: 0H and 48H reverting cells. Blue dots represent significantly up-regulated genes in 0H cells and purple dots represent significantly up-regulated genes in 48H reverting cells. Panel E: 48H reverting and 48H differentiating cells. Purple dots represent significantly up-regulated genes in 48H reverting cells and red dots represent significantly up-regulated genes in 48H differentiating cells.

Interestingly, when comparing 0H with 48H reverting cells, we saw only one gene that was significantly differentially expressed just above the threshold (FIGURE 3: panel D), the *RSFR* (RNase Super Family Related) gene, that is highly expressed in precursor cells from chicken bone marrow (26). Furthermore, when comparing 48H reverting with 48H differentiating cells, we found hemoglobin related genes up in the differentiating cells and *LDHA* and *ID2* up in reverting cells (FIGURE 3: panel E).

We more closely investigated gene expression distributions within the different conditions to see how gene expression patterns would evolve during the reversion process (FIGURE 4). We selected 8 genes differentially expressed and which expression increases or decreases during the differentiation process. *HBA1, HBBA, HBAD* (different hemoglobin subunits) and *FECH* (Ferrochelatase) are involved in hemoglobin and heme pathways and are more expressed by differentiating cells while *LDHA, ID2, CSTA* (cystatin A1) and *CRIP1* (Cysteine-rich intestinal protein1) are more expressed by self-renewing undifferentiated cells. We plotted and compared their distribution between the 4 conditions. For the genes involved in differentiation, we see a gradual shift in the distributions towards a higher level of expression as cells get more differentiated (FIGURE 4: Panel A - D) and we see the opposite shift for genes involved in proliferation (FIGURE 4: Panel E - H). In all cases, the 48H reverting cell expression patterns for those genes shifted back to patterns closer to the 0H cells. At the time of observation and especially for genes up in differentiation, the 48H reverting cell expression patterns are not completely similar to those of 0H cells. This was further confirmed by using a dedicated statistical tool, Sparse PLS (see below).

**Figure 4:**
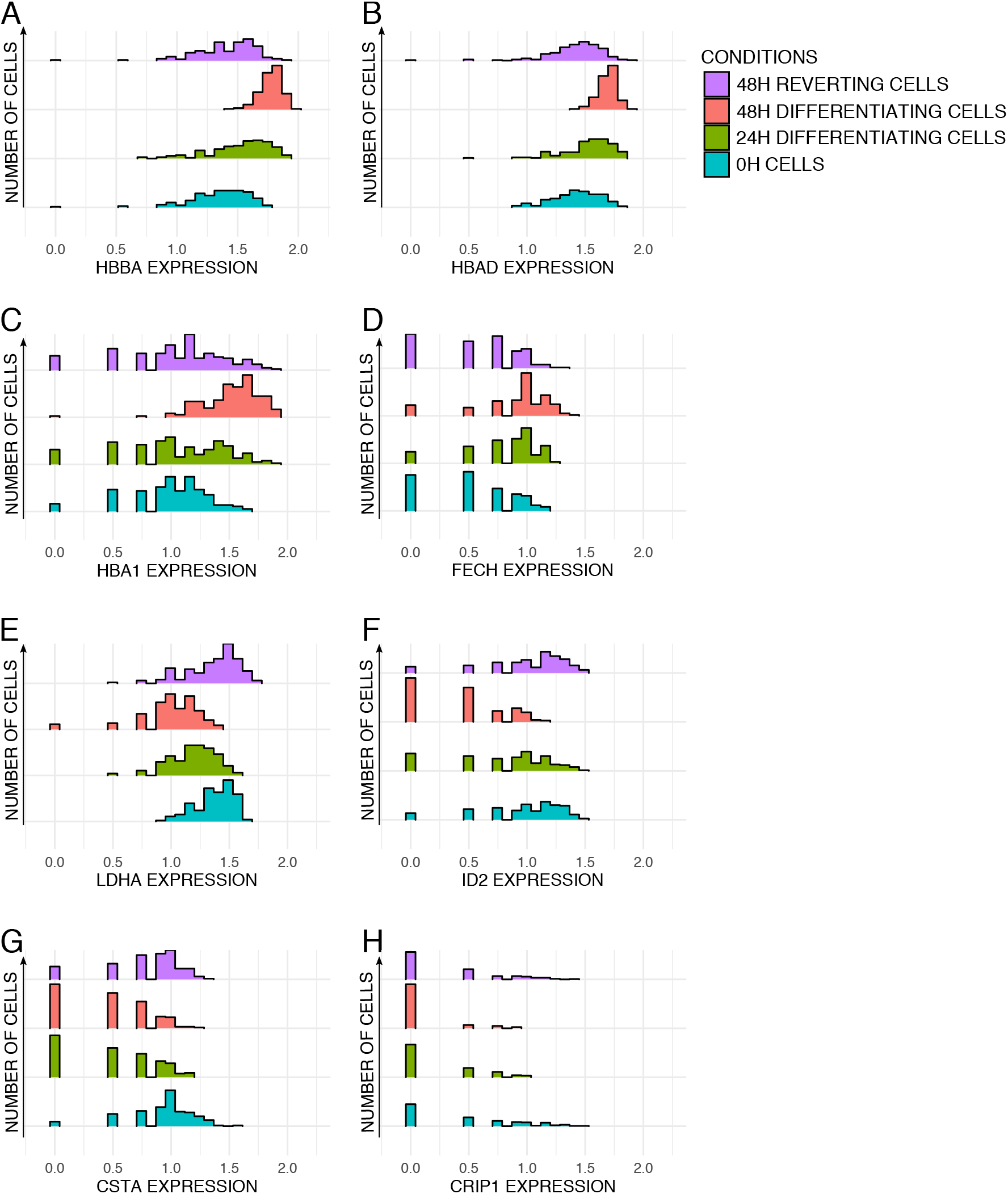
Comparison of gene expression pattern distributions between cells at four experimental time-points (0H, 24H, 48H differentiating and 48H reverting cells). X scale represents log1p of gene expression from scRNAseq data. Color legend: undifferentiated cells (0H in blue), 24H differentiating cells (in green), 48H differentiating cells (in red) and 48H reverting cells (in purple). Histograms of gene expression distribution for *HBBA* (Panel A), for *HBAD* (Panel B), for *HBA1* (Panel C), for *FECH* (Panel D), for *LDHA* (Panel E), for *ID2* (Panel F), for *CSTA* (Panel G) and for *CRIP1* (Panel H).

To go further on gene distribution comparisons we computed Wasserstein distances, a geometric distances well suited for comparing multimodal distributions, for each 2000 genes of the scRNAseq dataset between each condition two by two. We then obtain 6 distributions of Wasserstein distance values. Finally, we computed the Gini index as a measure of statistical dispersion in each distribution (the higher the Gini index is, the higher inequality among the values). We performed 100 bootstraps and compared the Gini values obtained (FIGURE 5A). Distribution of Wasserstein distances between 0H cells and 48H reverting cells had the smallest average Gini index among all distributions (Figure 5B). This result points towards a closer global transcriptional state between 48H reverting cells and 0H cells.

**Figure 5:**
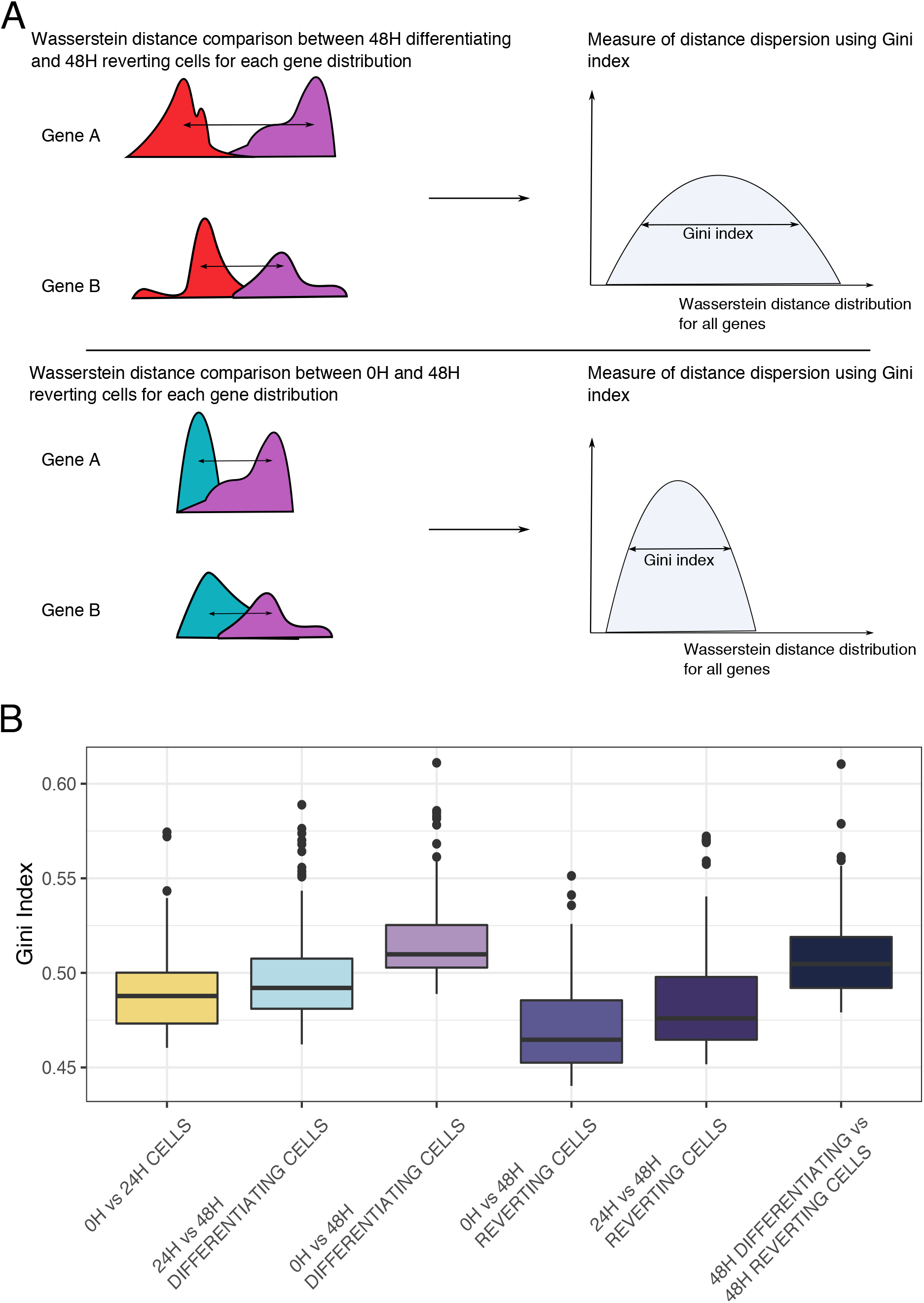
Comparison of dispersion of gene distribution between cell populations. Panel A: Experiment design to compare gene distributions between the 4 biological conditions. Wasserstein distance is computed for each gene between pair of conditions, then dispersion of all gene distributions is calculated using Gini index. Panel B: Plot of Gini index values of Wasserstein distance distributions between conditions in pairs computed for each of the 2000 genes from scRNAseq data bootstraped 100 times.

### 48H reverting cells retain molecular traces of a commitment into differentiation

To further characterize the molecular changes that persisted after reversion, we sought to identify predictive genes that discriminate the most the 48H reverting cells and the undifferentiated cells. We performed logistic regression combined with dimension reduction (Partial Least Square (20)) between 48H reverting cells and 0H cells and retained common most discriminating genes between scRT-qPCR and scRNAseq datasets. Interestingly, our results showed that only 3 common genes discriminate between the two cell groups: *HBBA, TBC1D7* and *HSP90AA1*, the expression of which is shown in FIGURE 6. *HBBA* is a subunit of the hemoglobin complex which carries oxygen, *TBC1D7* is presumed to have a role in regulating cell growth and differentiation (27) and *HSP90AA1* codes for an isoform of the HSP90 protein chaperone, which its specific transcription is known to be induced in response to insulin (28). Looking closely, the 48H reverting cells have an intermediate expression level between differentiating cells and undifferentiated cells for the three predicted genes. The offset observed could be due to a longer duration of mRNA half-life at 24H of differentiation. We had previously performed a quantification of mRNA half-life during avian erythrocyte differentiation ((29) Supplementary 2). We focused on mRNA half-life at 24H for those three genes. *TBC1D7* and *HSP90AA1* have a relatively short half-life as opposed to HBBA. Other genes analyzed whose expression increases during differentiation, such as *DPP7*, *TPP1* or *RPL22L1* have also a long half-life duration mRNAs, but only *HBBA* was identified in our statistical analysis as discriminating between undifferentiated and 48H reverting cells. These results confirmed that the 48H reverting cells display a gene expression pattern very close to those of 0H cells while still retaining traces of their engagement into the differentiation process independently of the mRNA half-life. The molecular process explaining such “lagging genes” will have to be explored.

**Figure 6:**
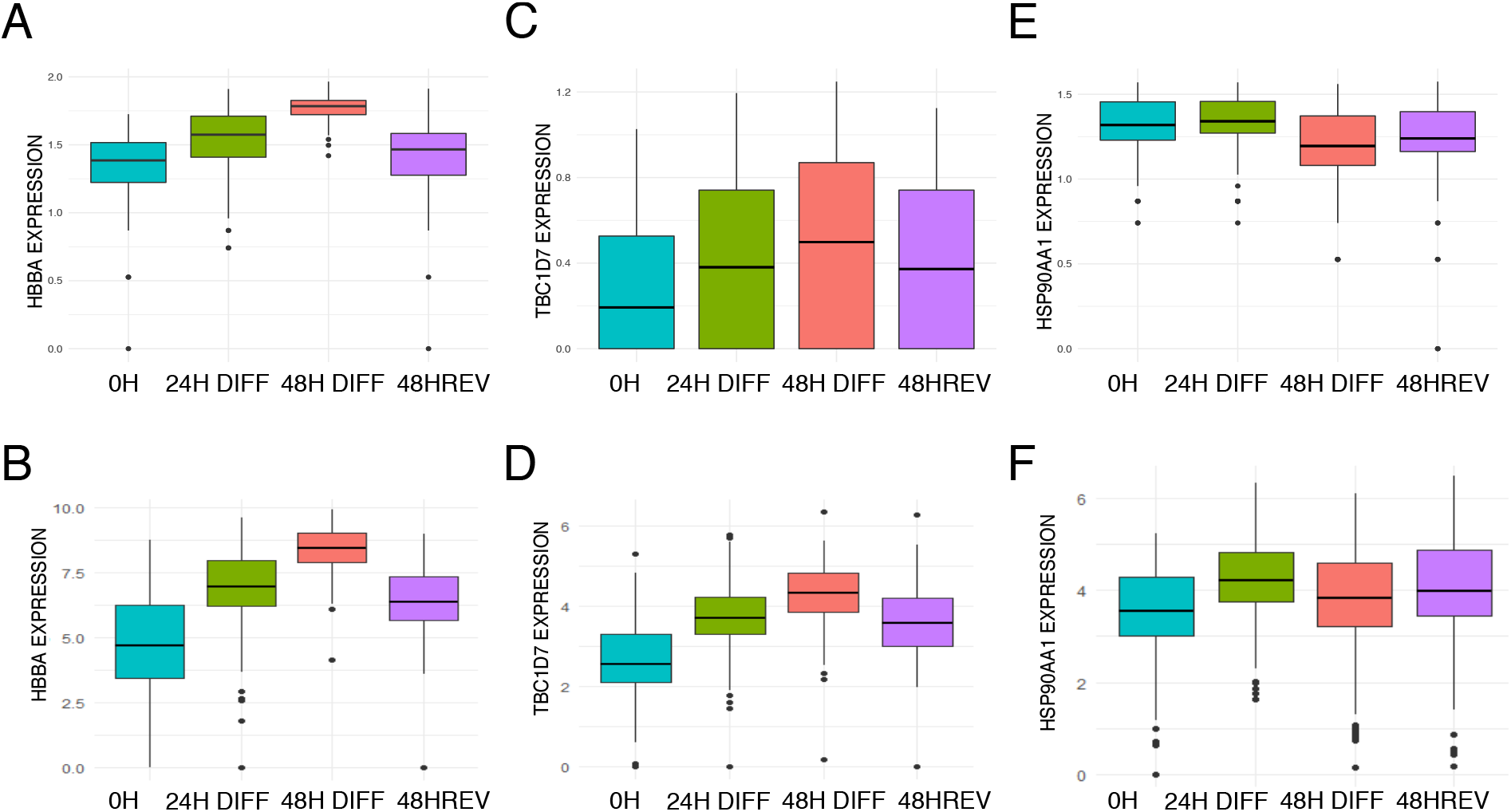
Boxplots with mean of expression levels of the 3 genes identified by Sparse PLS as discriminating genes between 48H reverting cells and 0H cells. Boxplots of *HBBA* expression level in log1p on scRNAseq data (panel A) and scRT-qPCR data (panel B) in the 4 biological conditions. Boxplots of *TBC1D7* expression level in log1p on scRNAseq data (panel C) and scRT-qPCR data (panel D) in the 4 biological conditions. Boxplots of *HSP90AA1* expression level in log1p on scRNAseq data (panel E) and scRT-qPCR data (panel F) in the 4 biological conditions.

### Cells are distributed as a continuum along the differentiation path

At that stage, two hypotheses could be made: 1. Either all cells have engaged into a differentiation process, and do molecularly revert to a self-renewal transcriptional state or 2. At 24H of differentiation two subpopulations coexist: one that is still undifferentiated and would give rise to the 48H reverting cells and a second more differentiated which would lead to the 48H differentiating population and die in the reversion experiment.

We hypothesized that the existence of two subpopulations at 24 hours should lead to a higher number of modes in the distribution of some genes at that time point. To test this hypothesis, we therefore estimated for each condition the most-likely number of modes for the probability distribution of each gene, as assessed through a Gamma mixture on scRNAseq (see material and methods). We found no significant difference in the number of modes observed between the 4 populations (Figure 7), which confirms that the cells collected from 24H do not show more multi-stability than the other groups, and are thus unlikely to be a mix of two populations.

**Figure 7:**
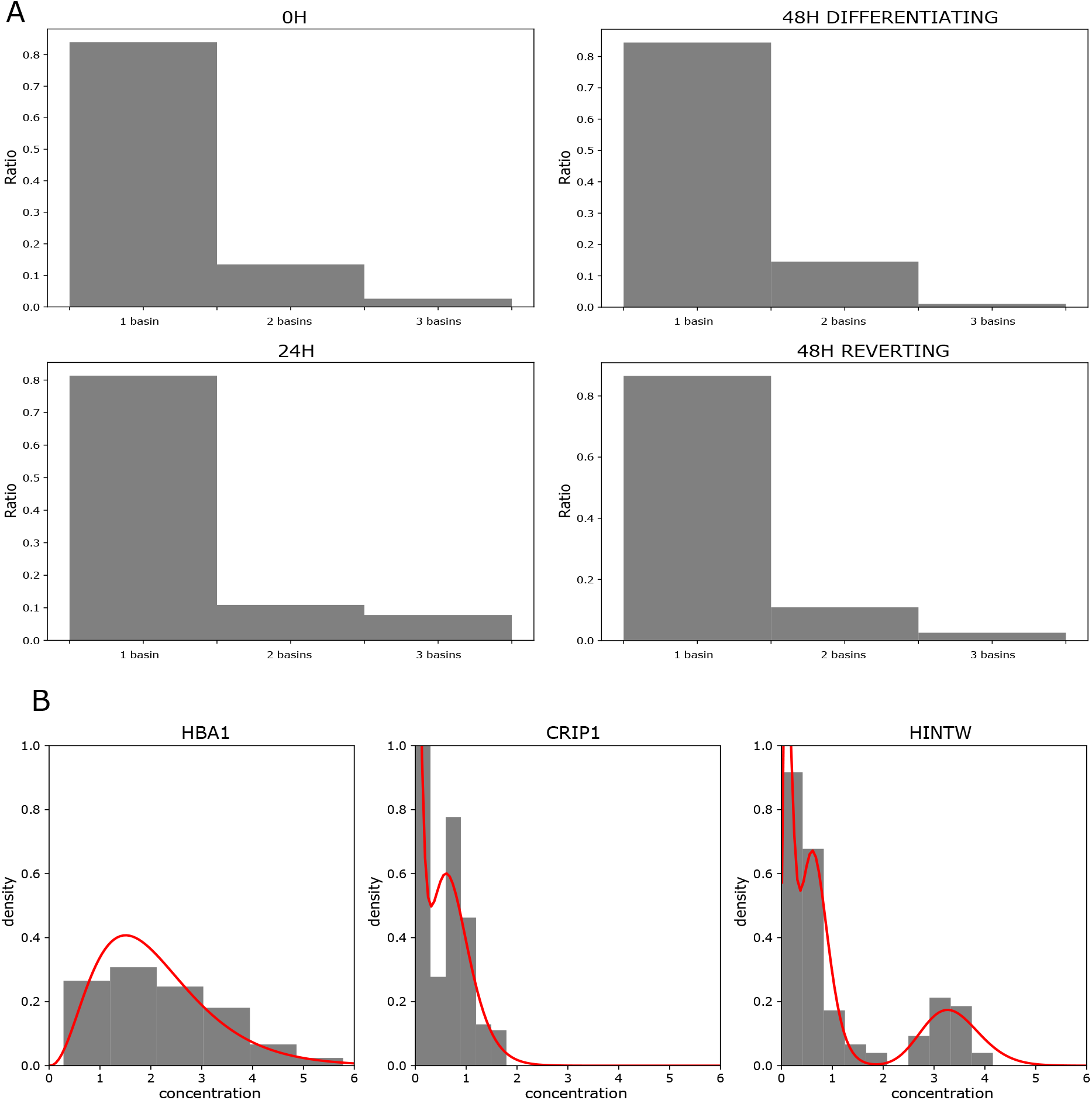
The repartition in the number of basins which have been detected for the 200 most variables genes from scRNAseq data, characterizing the level of multi-stability which is observed for each condition. Panel A: Repartition of the number of modes for each biological condition. Panel B: Examples of genes which distribution fits 1 basin (left), 2 basins (middle) or 3 basins (right).

The second hypothesis would also imply that in the 24H population the cells engaged too far in the differentiation process would die a short time after media was changed, while only the undifferentiated ones would survive. We then measured the viability rate during the kinetics and found no difference in viability between the conditions and especially between the 24H differentiation and the 48H reversion conditions (Supplementary 3).

Finally, the second hypothesis would also imply that the reversion cells are simply cells that have not entered the differentiation process. It would therefore be at odds with the evidence that the 48H reverting cells display slightly different pattern of gene expression as 0H cells, but do retain traces of their engagement into the differentiation process (see upper).

Those results strongly suggest that the 24H cell population is not composed of two coexisting subpopulations of cells and that 48H reverting cells enter differentiation before going back to a transcriptomic state close to 0H cells.

## Discussion

In the present study, we couple two different single-cell transcriptomic techniques and state-of-the-art statistical approaches to demonstrate the fate reversibility of avian erythrocyte progenitors induced to differentiate for 24 hours.

Our results show a very close proximity of reverting and undifferentiated cell transcriptomes. Indeed, statistical comparison of cell distributions showed no significant difference between 0H and 48H reverting cells while, as expected, significant changes in gene expression accompanied the differentiation sequence. The analysis of gene expression distribution patterns of the 48H reverting cells confirmed a switch toward the 0H cells gene expression profiles. First, DE analysis of scRNAseq data showed only one gene significantly differentially expressed between the two conditions. Second, Wasserstein distance analysis revealed closer distances between 48H reverting and 0H cells than to any other group of cells. Third, Sparse PLS analysis indicated that the expression level of only three gene, *HBBA, TBC1D7* and *HSP90AA1*, was predictive of the 48H reverting and undifferentiated cells. Interestingly the persistence of those three genes in 48H reverting cells does not seem to be caused by mRNA half-life duration.

All of our results therefore favor the hypothesis that a vast majority of the 48H reverting cells responded to differentiation induction by modifying their gene expression profiles but then returned to the self-renewal transcriptional state.

One must note that this would not be the sole example of large scale transcriptomic changes on (relatively) short time scales (18,30). The question as to whether such large-scale transcriptome changes are accompanied, or not, by (reversible) large scale epigenetic changes remains an open question for future studies.

It has been described in the literature that during cellular decision making, the cell state is maintained by dynamic interactions between positive and negative regulatory molecules (31) within the frame of a Gene Regulatory Network (GRN). These interactions can be repurposed by changing the stoichiometry of ubiquitous and specific regulatory molecules and factors (11,32). In our study, the analysis of gene expression patterns during the reversion process confirmed that the determination of the fate of erythrocyte progenitors is directed by the constraints of the dynamics of the GRN, influenced by signals emitted by changing conditions of the environment surrounding the cells. In the absence of differentiation signals (or in the presence of self-renewal inducing signals), there is no ratchet in place that would prevent (at least at early stages in our case) the system to return back to its original quasi-steady state. This is in excellent agreement with the previous demonstration that there is a duration threshold for some GRN under which the systems can return back to its original state (33). This was proposed to allow cells to discriminate between *bona fide* signals and random noise in their environment, and could represent a physiological system for finely tuning the *in vivo* production of red blood cells while preserving the pool of progenitors. We recently proposed a methodology for inferring the GRN underlying T2EC differentiation (29). For that we kept *in silico* cells under constant differentiation stimulus. It would be of interest to see if the inferred GRN would be able to revert, up to a certain point where no “spontaneous” return is possible (19), to its original state. This would be a very strong constraint to impose and should severely limit the number of putative GRN able to reproduce experimental data and thus approaching the most accurate network.

Taken together, our results point towards a physiological plasticity and reversibility with respect to erythrocyte decision-making. It is also reminiscent of the plasticity observed in Cancer Stem Cells that might not be specific to tumour cells (34). In terms of epigenetic landscape, our work implies that instead of a continuous gradient that the cells will roll down as in the classical Waddington’s depiction (8), they may go through an unstable state and may, sometimes, roll upwards over a bump in the landscape (35). Thus, differentiation should be more appropriately described as cells moving from well to well, that is, from one metastable state (36–38) to another one (Figure 8). This view abides by the multi-stability framework where a complex quasi-potential landscape aims at describing both normal and pathological differentiation processes (39,40), and exemplifies the fact that “commitment (is) a dynamical property of the landscape” (41). It is important at this stage to remember that Waddington himself was aware that his drawing was but a simplification. Adapting and refining this landscape should not be considered as departing from his views. Such a non-monotonous landscape has been proposed to account for the depiction of regeneration in adult tissues (42), and is consistent with previously proposed dynamical principles of cell fate restriction (10). It is in excellent accordance with the recent depiction that cells can “climb uphill on Waddington’s epigenetic landscape” during cranial neural crest cells development (15), and would also be more relevant to account for the « hesitant » behavior of human CD34 stem cells in vitro (43) than a straight slope. It is beyond the scope of this discussion to go into more details, but a cell “climbing uphill” should be seen as equivalent as “the landscape bending into a new valley”.

**Figure 8:**
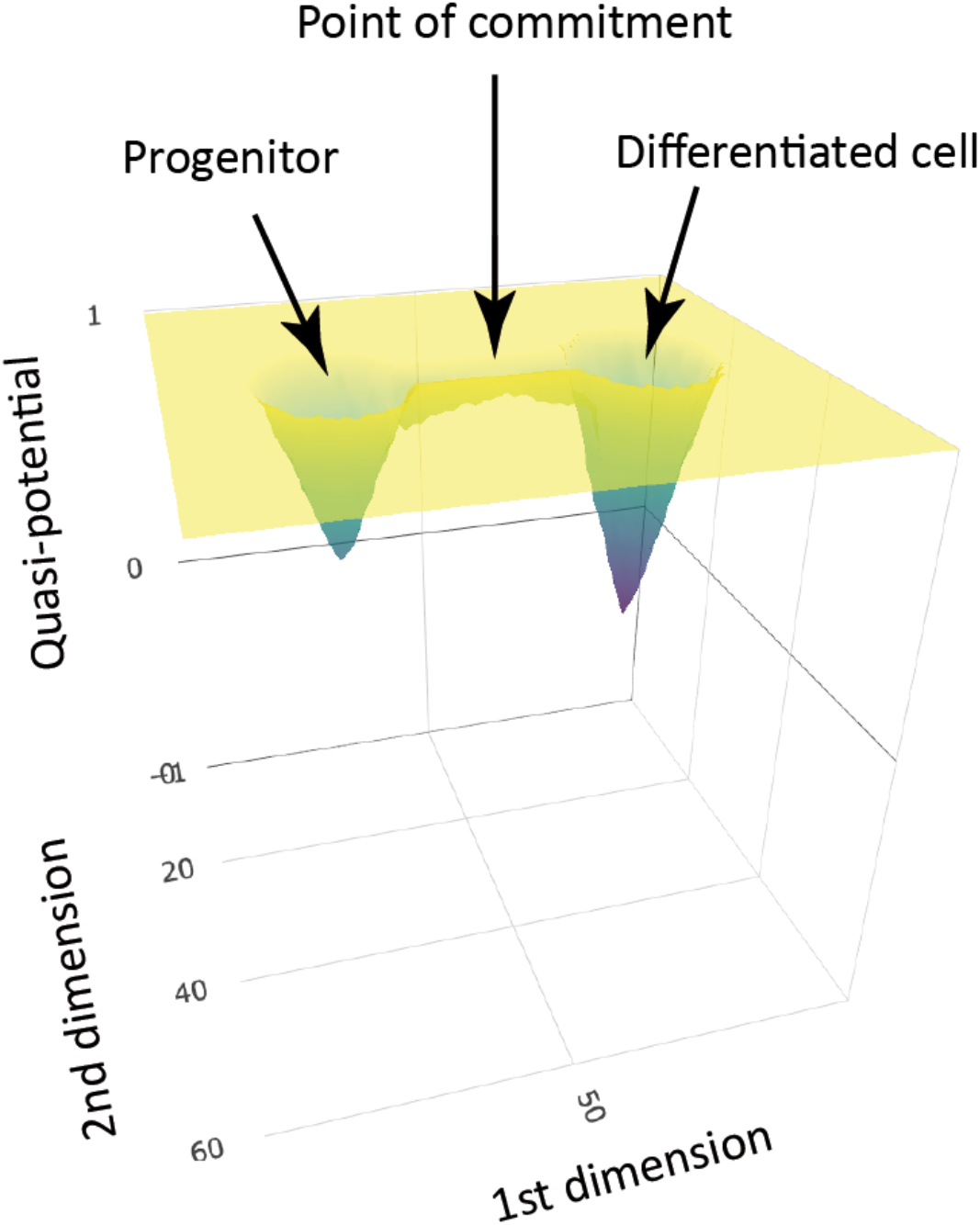
A quasi-potential well depictions of the erythroid differentiation process. While the cells have not escaped the zone of influence of the progenitor attractor (i.e. when they have not passed the point of commitment, *aka* the point of no return (19)) the removal of the environmental influences results in their relaxing back to their original attractor state.

In conclusion, our work has provided a detailed molecular characterization of the probabilistic nature of erythrocyte cell fate determination, influenced by the constraints of the underlying Gene Regulatory Network dynamics, and driven by environmental influences. These new insights into the process of cell reversion could also lead to significant improvements of the executable GRN inference scheme (29).

## Material and methods

### Cellular biology

T2EC were extracted from bone marrow of 19-d-old SPAFAS white leghorn chicken’s embryos (INRA, Tours, France). Cells were grown in self-renewal in LM1 medium (α-MEM, 10% Foetal bovine serum (FBS), 1 mM HEPES, 100 nM β-mercaptoethanol, 100 U/ mL penicillin and streptomycin, 5 ng/mL TGF-α, 1 ng/mL TGF-β and 1 mM dexamethasone) as previously described (44).

Differentiation was induced by removing LM1 medium and placing the cells into DM17 medium (α-MEM, 10% foetal bovine serum (FBS), 1 mM Hepes, 100 nM β-mercaptoethanol, 100 U/mL penicillin and streptomycin, 10 ng/mL insulin and 5% anemic chicken serum (ACS; (45)).

Differentiation kinetics were achieved by collecting a sub fraction of the cells at different times after induction of differentiation (0H and 24H). After 24H, DM17 medium was removed and half of the cells were placed back into LM1 medium while the other half was kept in DM17 medium to achieve 48H revertion and 48H differentiation time points respectively (FIGURE 1).

Cell population mortality was assessed by counting dead and living cells from the different time points and conditions after Trypan blue staining and using a Malassez cell.

### Single-cell sorting

Single-cells were sorted using a FACS Aria IIμ, BD. We collected around 200 cells per time point (8 plates) for each experiment (scRT-qPCR and scRNAseq, see below).

### Single-cell RT-qPCR analysis

All the manipulations related to the high-throughput scRT-qPCR experiments in microfluidics were performed according to the protocol recommended by the Fluidigm company (PN 68000088 K1, p.157-172). All steps from single-cell isolation to scRT-qPCR, genes selection, data generation and cleaning are described in detail in (19). Expression matrix was log1p transformed before subsequent analysis.

### Single-cell RNAseq

scRNAseq was performed using an adapted version of MARSseq protocol (Massively parallel single-cell RNA sequencing) (22). Unless specified, all indicated concentrations correspond to final concentrations.

Individual cells were sorted into 96-well plates containing 4μL of lysis buffer and index RT primers (0,2% Triton (Sigma Aldrich), 0,4 U/μL RNaseOUT (Thermofisher Scientific), 400nM RT_primers (Sigma Aldrich)). Index RT_primers (Table 1) contain oligo-dT chain to capture mRNA, a T7 RNA polymerase promoter for further *in vitro* transcription (IVT), unique cell barcodes for subsequent de-multiplexing and unique molecular identifiers (UMIs) for PCR bias deduplication. After cell sorting, plates were immediately centrifuged and frozen on dry ice before storage at −80°C until reverse transcription (RT) was performed. The plates were put at 72°C for 3 minutes for denaturation. 4μL of RT mix were added in each well (2mM dNTP (Thermo scientific), 20mM DTT, 2X First stranded buffer, 5 U/μL Superscript III RT enzyme (Superscript III RT enzyme kit Thermo scientific), 10% (W/V) PEG 8000 (Sigma Aldrich)). ERCC RNA spike-in (Thermo Scientific) were diluted into the RT mix (dilution 5×10^-7^). The plates were then transferred into a thermocycler (program: 42°C-2min, 50°C-50min, 85°C-5min, 4°C hold).

**Table 1:** P-values output of multivariate two tests between pair of conditions compared.

|  | 0H vs 48H reverting | 0H vs 48H differentiating | 48H reverting vs 48H differentiating |
| --- | --- | --- | --- |
| p-value | 1.00 | 0 | 0.00000000369 |

After reverse transcription, samples were pooled by plate and ExonucleaseI (NEB) digestion was performed, followed by 1,2X AMpure beads purification (Beckman Coulter). Samples were eluted in 10mM Tris-HCl, pH=7,5. Second strand cDNA synthesis was performed with 1X SSS buffer and SSS enzyme (NebNext mRNA second strand synthesis kit NEB; thermocycler program: 16°C-150min, 65°C-20min, 4°C hold). Resulting double strand cDNA were linearly amplified by IVT overnight (10mM ATP, 10mM GTP, 10mM UTP, 10mM GTP, 1X reaction buffer, 1/10 T7 RNA polymerase mix (HighScribe T7 High Yield RNA synthesis NEB)) at 37°C. IVT products were purified with 1,3X Ampure beads and eluted with 10mM Tris-HCl, 0,1mM EDTA. Amplified RNAs were fragmented (1X RNA fragmentation buffer (RNA fragmentation reagents Invitrogen)) at 70°C for 3 min. The fragmentation reaction was stopped with 34μL of STOP mix (0,3X Stop solution (RNA fragmentation reagents Invitrogen), TE buffer 1X (10mM Tris, 1mM EDTA, pH 8 - Invitrogen) and 0,7X AMpure beads to procede with sample purification). Differing from original MARSseq protocol, instead of ligation, a second RT was done to incorporate P5N6 primers (Table 2) containing random hexamers and specific barcodes to distinguish the different plates (5mM DTT, 500μM dNTP, 10μM P5N6_XXXX, 1X First stranded buffer, 10U/μL Superscript III RT enzyme, 2U/μL RNaseOUT; thermocycler program: 25°C 5min, 55°C 20min, 70°C 15min, 4°C hold). The cDNAs were then purified with 1,2x AMpure beads. Illumina primers (Table 1) were added by PCR (0,5 μM Mix primer P5.rd1/P7.Rd2, 1X KAPA Hifi HotStart PCR Mix (Kapa Biosystem); thermocycler program: 95°C 3min, 12 times [98°C 20sec, 57°C 30sec, 72°C 40sec], 72°C 5min, 4°C hold), and PCR products were purified with 0,7x AMpure beads and eluted in 15μL. Libraries were sequenced on a Next500 sequencer (Illumina) with a custom paired-end protocol to avoid a decrease of sequencing quality on read1 due to high number of T added during polyA reading (130pb on read1 and 20pb on read2). We aimed for a depth of 200 000 raw reads per cell.

**Table 2:**
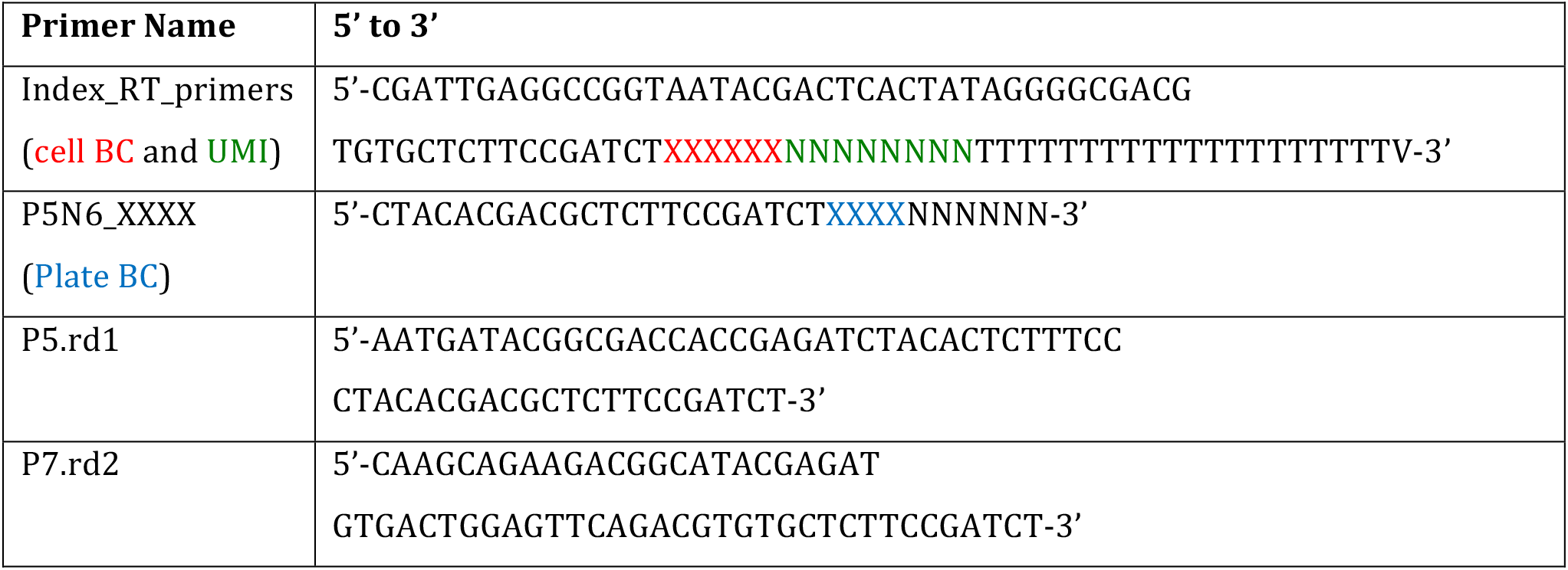
List and sequences of primers used for scRNAseq libraries construction.

### Bio-informatic pipeline

Fastq files were pre-processed through a bio-informatic pipeline developed in the team on the Nextflow platform (46). Briefly, the first step removed Illumina adaptors. The second step de-multiplexed the sequences according to their plate barcodes. Then, all sequences containing at least 4T following cell barcode and UMI were kept. Using UMItools whitelist, the cell barcodes and UMI were extracted from the reads. The sequences were then mapped on the reference transcriptome (Gallus GallusGRCG6A.95 from Ensembl) and UMI were counted. Finally, a count matrix was generated for each plate.

### Data filtering, normalization and analysis

All analysis were carried out using R software (4.0.5;(47)). Matrixes from the eight plates were pooled together. Cells were filtered based on several criteria: reads number, genes number, counts number and ERCC content. For each criteria the cut off values were determined based on SCONE (48) pipeline and were calculated as follows:

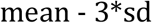

We selected genes present in at least two cells. Filtered matrix was then normalized using SCTransform from Seurat package (49) and we corrected for batch effect, time effect and sequencing depth effect. Expression matrix was finally log1p transformed. Variable genes were identified using FindVariableFeatures from Seurat, vst method (50). Based on visualization of genes variance, we retained the 2000 most variable features. Differentially expressed genes were identified using FindMarkerGenes function from Seurat (50). Analysis was by pairwise comparisons between conditions, genes with log fold change >=0,5 and adjusted p-value <0,05 were kept as significant. More information on QC filtering are given in Supplementary 4.

### Statistical analysis

All statistical analyses were performed using the R software (4.0.5; (47)). Dimensionality reduction and visualization were performed using UMAP (51). Adaptive Sparse PLS for Logistic Regression was performed using the plsgenomics package (20). For this analysis, scRT-qPCR data were scaled. Sparse PLS is a supervised statistical analysis that allow to predict the most discriminant variables between two groups. Wasserstein distance computation was done using the Transport R package (52), and was accomplished for each gene of the scRNAseq dataset.

Gini indexes were calculated using the Ineq R package on Wasserstein distance distributions (53).

Bootstraping was done using sample_frac function from Dplyr R package(54).

### Estimation of multi-stability levels

For estimating the level of multi-stability in the data, we considered that the probability distribution of each gene can be approximated by a Gamma distribution, or a mixture of Gamma distributions, since they are known to describe continuous single-cell data accurately (55). More precisely, we parameterized the distribution of a gene *i* by:

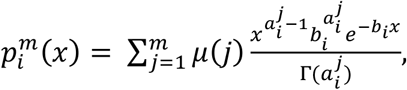

Where Γ denotes the Gamma function: 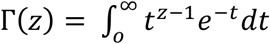. Note that only the parameters 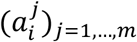 depend on the mixture component *j*: this is related to the distribution arising from the well-established two-states model of gene expression (56), when only the frequency of mRNA bursts is regulated, as described in (57).

For every condition, we constructed 10 training sets consisting of 80% of the cells in the population (randomly-chosen), and we estimated the parameters 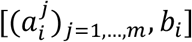 with a MCMC algorithm for the numbers of mixture components *m* = 1,2, 3 successively. We then considered that the optimal number of components for gene *i* was the one which minimized the average BIC score estimated on the 10 corresponding test sets.

### Multivariate two-sample test

Samples were compared using a multivariate two-sample test based on the 2000 most variable genes. We suppose that the normalized gene expression X_1_ and X_2_ of two conditions (0H vs 48H reversion, 0H vs 48H differentiation, 48H reversion vs 48H differentiation), follow a multivariate Gaussian distribution 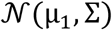 and 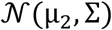 respectively, and we denote by *n* = *n*_1_ + *n*_2_ the total number of cells. Then we test the null hypothesis *H*_0_: *μ*_1_ = *μ*_2_ using the generalized Hotelling’s *T*^2^ test (23). The data being high dimensional (*p* > *n*), the between-gene pooled covariance matrix is not invertible, and is replaced by its Moore-Penrose inverse. In this setting the asymptotic distribution of the generalized Hotelling statistics is χ^2^ (*n* – 2). The p-values were adjusted according to the Benjamini-Hochberg correction (58). Analysis was performed using the fdahotelling R package (59).

## Code and data availability

Pipelines and analysis scripts are available at https://gitbio.enslyon.fr/LBMC/sbdm/mars_seq. scRT-qPCR data are available at https://osf.io/upw8d/. scRNAseq data are available at http://www.ncbi.nlm.nih.gov/bioproject/802343 BioProject ID PRJNA802343.

## Acknowledgements

We gratefully thank all members of SBDM team and particularly Gerard Benoit for very fruitfull discussions, suggestions and commentaries on our project. We also thank Ghislain Durif for his great technical support during PLS computation and G. Yvert for helpful comments about the manuscript. We thank the computational center of IN2P3 (Villeurbanne/France) and Pôle Scientifique de Modélisation Numérique (PSMN, Ecole Normale Supérieure de Lyon) where computations were performed. We aknowledge the contribution of the AniRA-Cytométrie core facility of SFR BioSciences (UAR3444/US8). We thank the BioSyL Federation and the LabEx Ecofect (ANR-11-LABX-0048) of the University of Lyon for inspiring scientific events.

