## supplementary for "Evidence for close molecular proximity between reverting and undifferentiated cells"

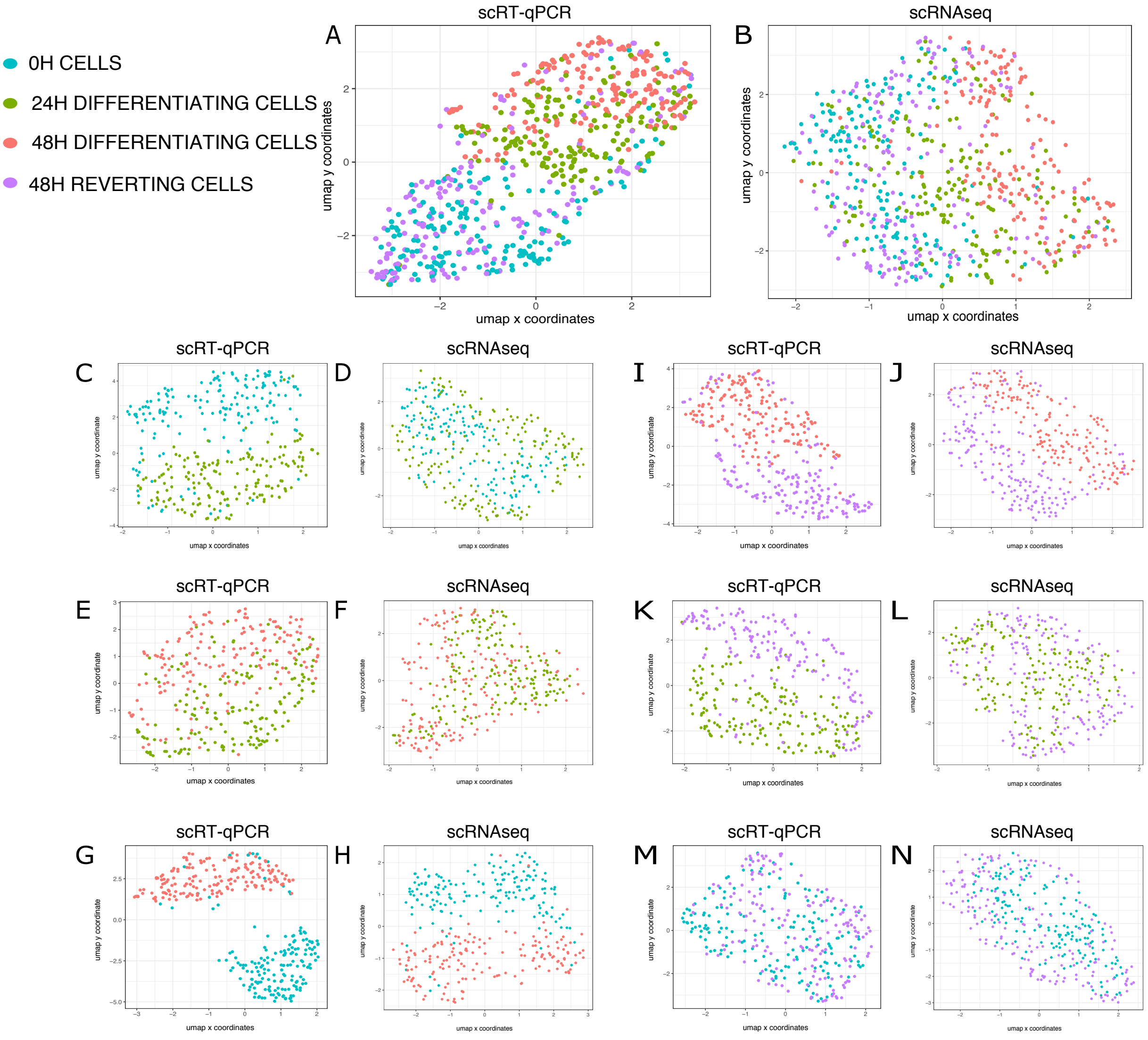


Supplementary 1: UMAP visualization of scRT-qPCR and scRNAseq data. All UMAPs are recalculated when projecting conditions two by two. 0H cells are displayed in blue, 24H cells in green, 48H differentiating cells in red and 48H reverting cells in purple.

Panels A, C, E, G, I, K and M: scRT-qPCR data

Panels B, D, F, H, J, L and N: scRNAseq data

Panels A and B: all 4 conditions

Panels C and D: 0H and 24H differentiating cells

Panels E and F: 24H and 48H differentiating cells

Panels G and H: 0H and 48H differentiating cells

Panels I and J: 48H differentiating and 48H reverting cells

Panels K and L: 24H and 48H reverting cells

Panels M and N: 0H and 48H reverting cells

| Gene Name | Ensembl_Gene_ID | mRNA half-life at 24h (hour) |
|---|---|---|
| HSP90AA1 | ENSGALG00000011351 | 3,47 |
| TBC1D7 | ENSGALG00000012731 | 3,69 |
| RPL22L1_1 | ENSGALG00000009312 | 16,64 |
| betaglobin | ENSGALG00000028273 | 16,64 |
| TPP1 | ENSGALG00000022706 | 16,64 |
| DPP7 | ENSGALG00000009105 | 16,70 |

Supplementary 2: Half-life of mRNA

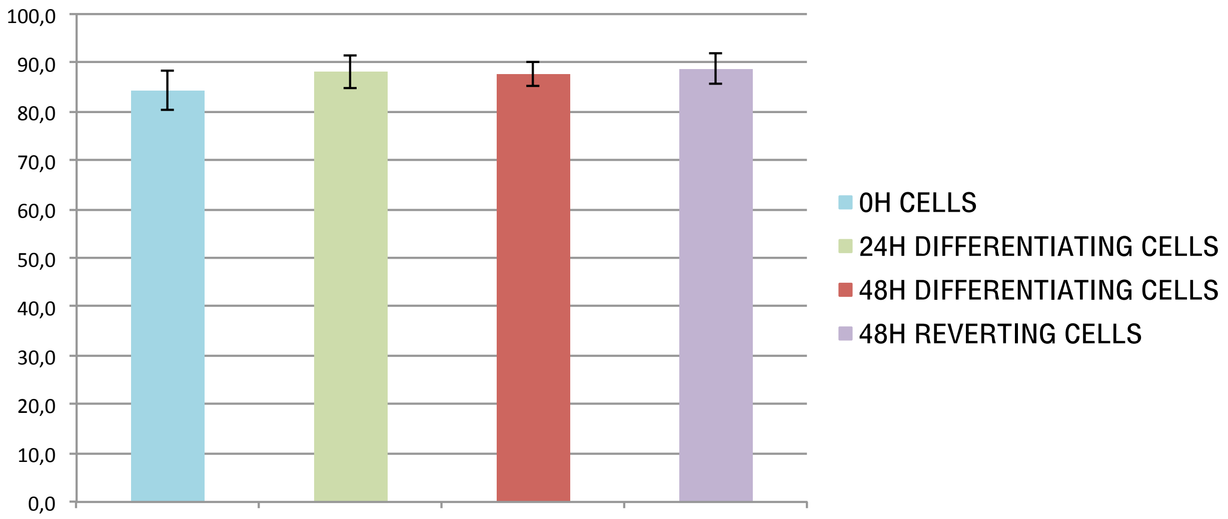


Supplementary 3: Histograms of viability rate during reversion and differentiation processes measured by Trypan blue staining and counting in a Malassez chamber. Means were compared using Student t-tests.

| QC step | Filter value | Number of cells remaining | Number of features remaining |
|---|---|---|---|
| **Number of sequencing reads (per cell)** | 80000 reads | 741 cells | / |
| **Percentage of mapped reads (per cell)** | 46% | 740 cells | / |
| **Percent of reads mapped to ERCC (per cell)** | <20% | 710 cells | / |
| **Number of detected genes (per cell)** | 233 genes | 710 cells | / |
| **Number of UMIs (per cell)** | 1584 UMIS | 710 cells | / |
| **Normalization** | / | 710 cells | 8682 genes |
| **Variable features selection** | Vst 2000 most variable genes | 710 cells | 2000 genes |

Supplementary 4: Summary table of scRNAseq data filtering steps and threshold values for each step. Threshold determination is explained in material and methods.
